# SnRK1.1 Coordinates Organ-Specific Growth–Defense Programs via Transcriptomic Rewiring in *Arabidopsis thaliana*

**DOI:** 10.1101/2025.04.25.650715

**Authors:** Tetiana Kalachova, Karel Müller, Jozef Lacek, Simon Pree, Anzhela Antonova, Oleksandra Bondarenko, Lenka Burketová, Katarzyna Retzer, Wolfram Weckwerth

## Abstract

SnRK1 (Sucrose non-fermenting-1-Related Kinase 1) is a master regulator of cellular energy homeostasis in plants, coordinating developmental and metabolic responses under environmental and internal stress conditions. Here, we demonstrate that its catalytic subunit, KIN10, orchestrates organ-specific growth–defense programs in *Arabidopsis thaliana* through transcriptomic rewiring. Using RNA-seq profiling, we reveal that *kin10* knockout mutants exhibit extensive transcriptional reprogramming in roots, particularly in pathways linked to signal perception, cell wall remodeling, and intracellular trafficking, which correlates with impaired root growth and reduced root hair elongation upon *Pseudomonas syringae* infection. In contrast, KIN10 overexpression (KIN10-OE) lines display constitutive defense activation in shoots, including elevated expression of genes associated with reactive oxygen species (ROS) production and salicylic acid (SA) signaling, leading to enhanced ROS accumulation and growth trade-offs. *kin10* roots show muted responses to biotic stimuli, while KIN10-OE shoots prioritize defense over growth. These findings position KIN10 as a critical integrator of energy status and immune signaling, enabling fine-tuned, tissue-specific responses essential for optimizing plant adaptation to dynamic environmental challenges.

## 1. Introduction

SnRK1 (Sucrose non-fermenting-1-Related Kinase 1) is a central regulator of cellular energy homeostasis, coordinating metabolic and developmental processes in response to environmental and internal cues (1–3). Its catalytic subunit, SnRK1.1/ KIN10/SnRK1α, has been identified previously as a crucial player in regulating growth, stress responses, and biotic interactions (1,4,5). Previous studies have highlighted the role of SnRK1 in regulating key processes, including meristem activity, root elongation, and root hair development (1–4,6). In particular, SnRK1’s activity in the root epidermis, where root hairs emerge, suggests its importance in integrating energy status with cell-specific responses (1).

Overexpression of KIN10 has been associated with reduced shoot growth, likely due to constitutively active stress responses that prioritize survival over growth (6). Conversely, kin10 mutants display impaired root development, including deregulated meristem activity and reduced root hair elongation (2). Root hair elongation, a key adaptation during interactions with both beneficial and pathogenic microbes, requires a tightly coordinated interplay between ROS production, hormonal signaling, and cellular reorganization (2). Pathogens, such as *Pseudomonas syringae* (Pst), exploit these processes to establish infections (7). However, the extent to which KIN10 activity modulates these pathways during pathogen challenge remains unclear. Our findings highlight the dual role of KIN10 as a coordinator of growth and defense, emphasizing its importance as a central signaling hub in plant adaptation.

Here, we show that KIN10 exerts distinct regulatory functions in roots and shoots, particularly in the context of microbial sensing and defense-related processes. Transcriptomic profiling revealed that roots of the established KIN10 knockout mutant (*kin10*) exhibit strong transcriptional reprogramming, especially in pathways related to signal perception, cell wall remodeling, intracellular trafficking, and kinase activity, all of which are essential for cell elongation and root hair development. These changes correlate with the impaired root responses upon infection, including numbness regarding root growth response and reduced root hair elongation, which demonstrates a weakened response to pathogen challenge.

In contrast, shoots of the KIN10 overexpression line (KIN10-OE) shows already without infection constitutive defense activation, including enhanced expression of genes involved in reactive oxygen species (ROS) production and salicylic acid signaling. These molecular responses align with elevated ROS accumulation and growth trade-offs typically associated with stress prioritization. Despite reduced developmental growth, KIN10-OE shoots demonstrate an enhanced defense capacity, suggesting that KIN10 modulates the balance between growth and immune activation in an organ-specific manner.

Together, our findings highlight KIN10 as a tissue-specific integrator of energy and immune signaling, essential for fine-tuning plant growth and defense in response to environmental cues.

## 2. Results

### 2.1. Antagonistic orchestration of root and shoot by KIN10

SnRK1 is a central regulator of cellular activity. Its catalytic subunit, KIN10, is essential for orchestrating metabolic and developmental processes, as well as enabling efficient responses to environmental signals, including interactions to biotic stimuli. To investigate the organ-specific functions of KIN10, we performed RNA-seq analysis comparing *kin10* and KIN10-OE to their respective wild-type backgrounds (*Col-0* and *Ler-0*). Differential expression analysis revealed that *kin10* roots exhibit substantial transcriptional reprogramming, with most differently expressed transcripts being downregulated, whereas KIN10-OE plants display more pronounced alterations in the shoot (Fig. 1A). Gene ontology (GO) enrichment analysis further demonstrated that *kin10* roots downregulate genes associated with signal perception, cell wall remodeling, and intracellular trafficking (Fig. 1B). Those pathways are critical for root elongation and defense responses. In contrast, KIN10-OE shoots showed strong upregulation of defense-related genes, particularly those involved in reactive oxygen species (ROS) production and salicylic acid (SA) signaling pathways. These transcriptomic patterns suggest that KIN10 acts as an organ-specific regulator, coordinating a fine balance between growth and immune activation depending on the tissue context.

**Figure 1:**
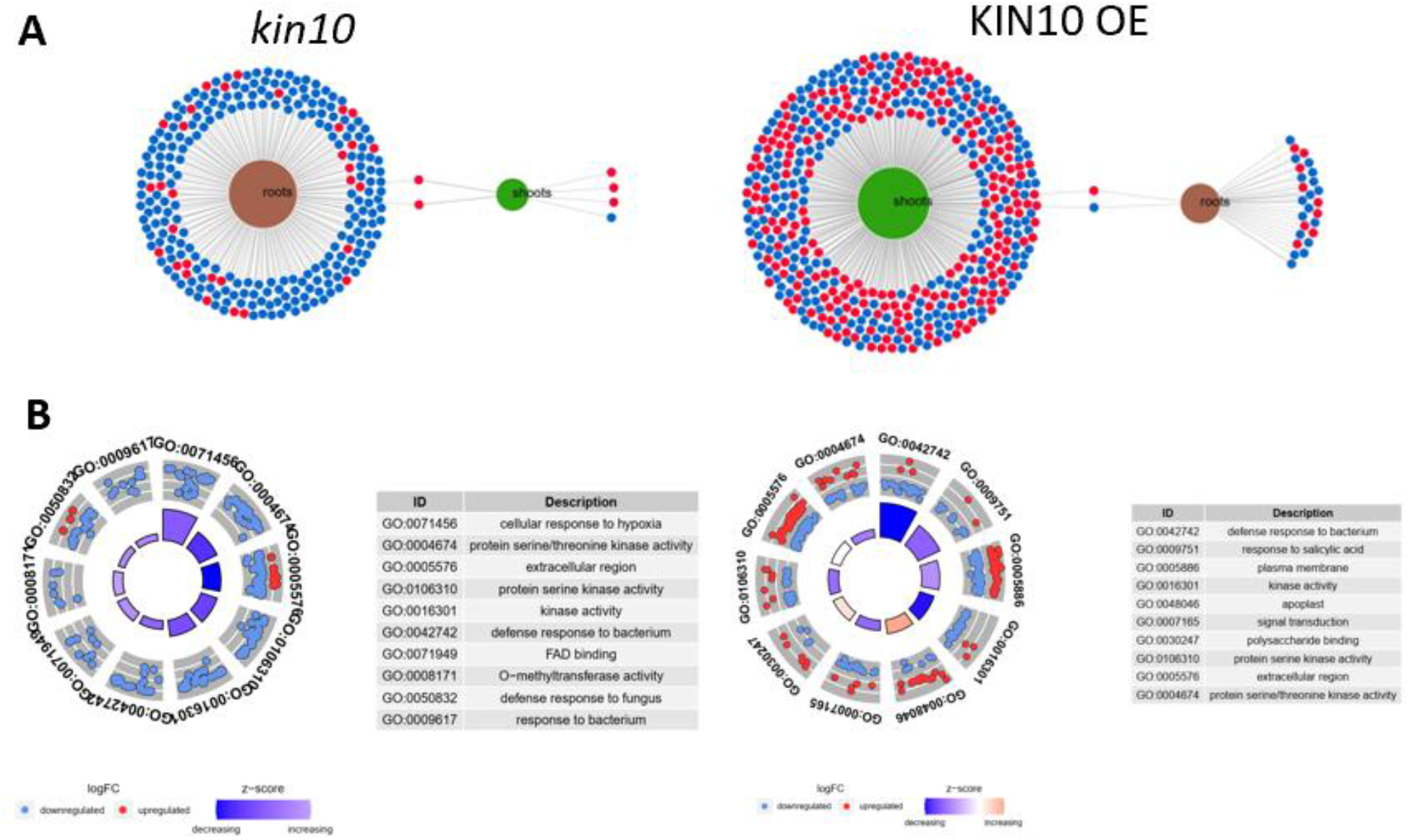
(A) Differentially expressed genes (DEGs) in roots (*kin10* vs. *Col-0*) and shoots (KIN10-OE vs. *Ler-0*) from RNA-seq data. (B) GO enrichment shows *kin10* roots downregulate genes related to signal perception, cell wall remodeling, and trafficking. KIN10-OE shoots upregulate defense-related genes including those involved in ROS production and SA signaling. Transcriptomic shifts suggest organ-specific regulation of growth-defense balance by KIN10. Overall, *kin10* show substantial transcriptional changes in roots compared to wild type plants, whereas KIN10-OE show more pronounced alterations in the shoot.

### 2.2 *kin10* roots show reduced responses to *Pseudomonas syringae (Pst)* infection

Transcriptomic profiling revealed that *kin10* roots exhibit strong transcriptional reprogramming. We performed different stress assays, and the most profound differences for root occurred afterwards *Pseudomonas syringae* (*Pst*) infection, as visible in the images in Figure 2 (Fig. 2B). *kin10* mutants displayed a muted response compared to wild-type plants, showing less primary root growth inhibition (Fig, 2A) by bacterial peptide flg22 and significantly reduced root hair formation and elongation in response to virulent strain of *Pst* (Fig. 2 C-E). In contrast, KIN10-OE lines exhibited enhanced root growth inhibition and disproportionately increased root hair elongation, both under basal conditions and infectes, but still showed a significant response upon pathogen challenge (Fig. 2 C-E), which might result from a constitutively alerted condition.

**Figure 2:**
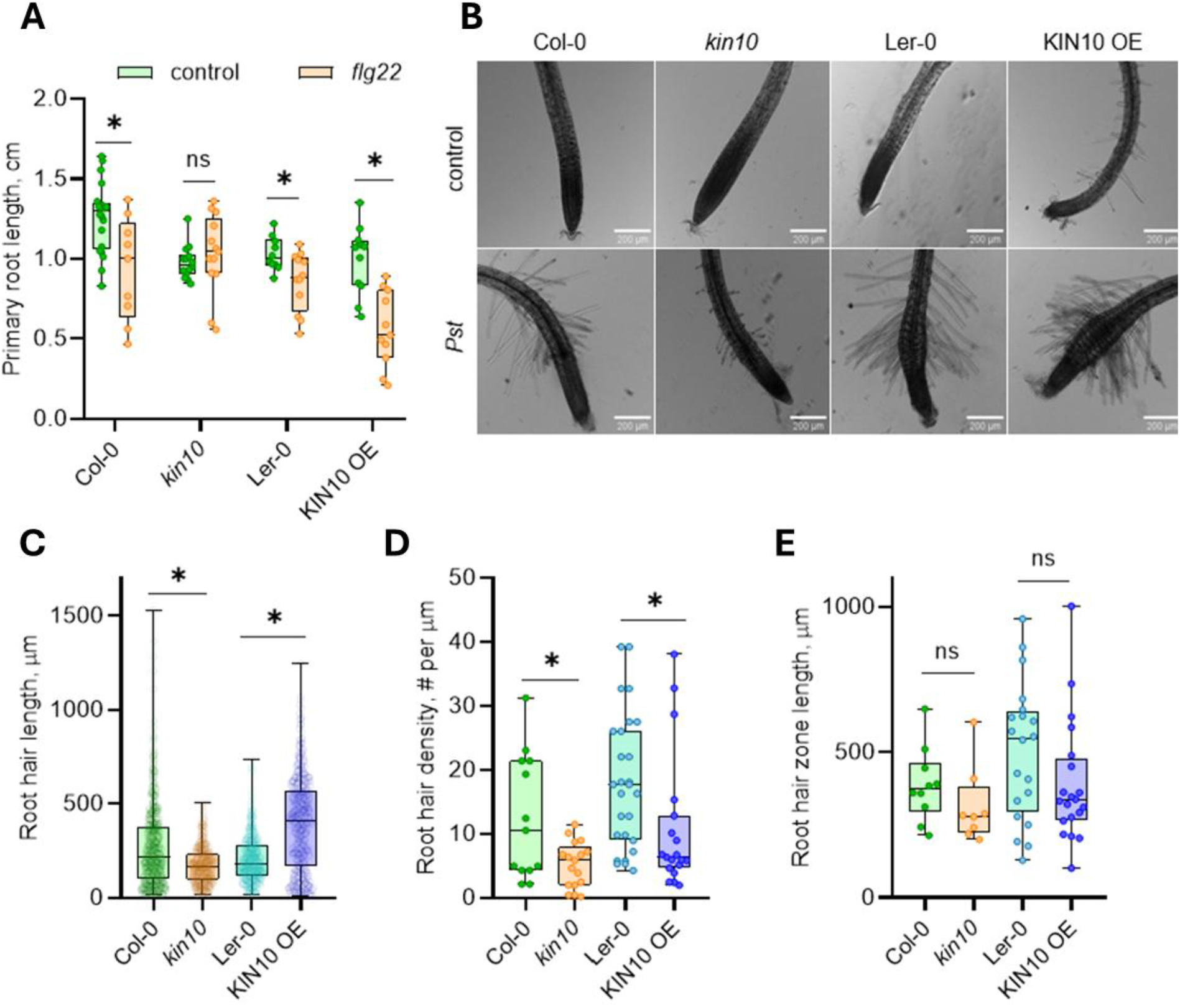
KIN10 modulates root growth inhibition and root hair elongation in response to biotic stimuli. (A) Root growth inhibition assay comparing primary root length of Col-0 and *kin10*, and Ler-0 and KIN10-OE seedlings grown on media with or without flg22. *kin10* did not exhibit attenuated root growth inhibition compared to wild-type or KIN10-OE plants. (B) Root hair elongation analysis showing *Col-0, kin10, Ler-0*, and KIN10-OE seedlings with and without *Pseudomonas syringae* (Pst) infection. *kin10* demonstrate reduced root hair formation and elongation, whereas KIN10-OE lines exhibit enhanced responses. (C) Quantification of root hair length (μm), (D) density (hairs/mm) and (E) length of root hair zone, illustrate that *kin10* display a muted response to *Pst* infection, characterized by less pronounced root growth inhibition and reduced root hair development compared to wild-type control.

**Figure 3.**
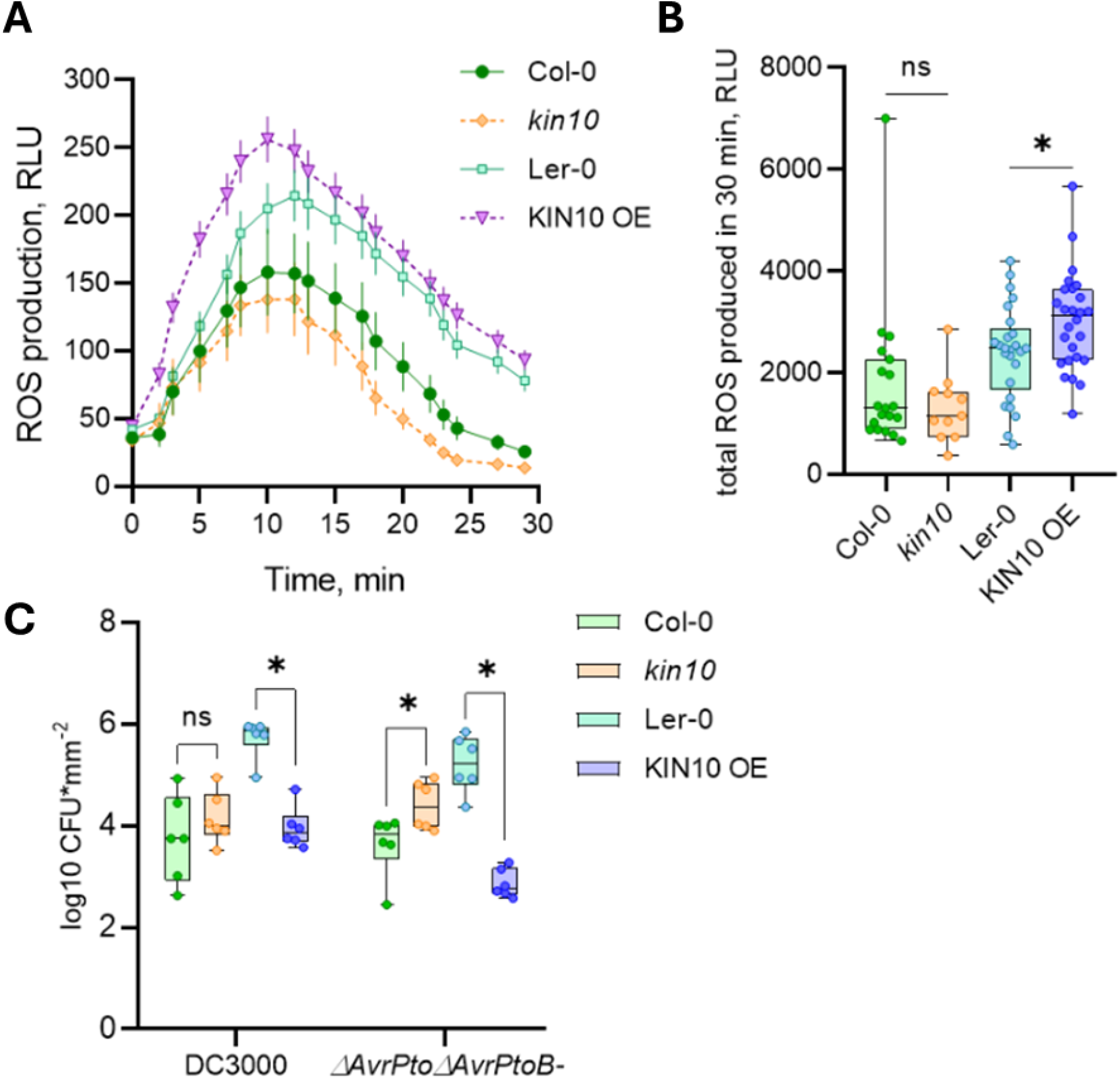
KIN10 overexpression enhances reactive oxygen species (ROS) burst in *Arabidopsis* leaves. (A) ROS production measured in *Col-0, kin10, Ler-0*, and KIN10-OE leaves following infection, using luminol-peroxidase assay. (B) Relative Luminescence Units (RLU) over 30 minutes show significantly elevated ROS production in KIN10-OE compared to controls. *kin10* shows wild-type-like ROS kinetics. (C) Responses to wild-type Pst DC3000 and effector-deficient mutated strain PtoΔvrPtoΔAvrPtoB-.

### 2.3. KIN10 OE in shoots shows constitutively activated defence response

KIN10-OE shoots transcriptomics showed among others the upregulation of genes involved in reactive oxygen species (ROS) production and salicylic acid signalling. These molecular responses align with elevated ROS accumulation and growth trade-offs typically associated with stress prioritization. Despite reduced developmental growth, KIN10-OE shoots demonstrate an enhanced defence capacity, suggesting that KIN10 modulates the balance between growth and immune activation in an organ-specific manner. To quantify the differences in ROS production we applied enzymatic analyses in leaves upon flg22 stimulation, which confirmed significant changes in ROS production depending on KIN10 expression levels. Rapid flg22-induced ROS production is mostly a result of activity of plasma membrane NADPH-oxidases RBOHD/F, which depends mostly on the receptor complex and enzyme presence. KIN10 deficiency or overexpression did not impact the dynamics of the process, but the overall amount of produced ROS during 30 min. was elevated in KIN10-OE line. For *kin10* we detected a wild type like curve of ROS production, whereas the overexpression line showed elevated ROS production.

## 3. Discussion

The results of this study underscore the critical role of KIN10 as a central regulator of SnRK1-mediated activity, influencing metabolic, developmental, and environmental response pathways. KIN10 activity is indispensable for orchestrating cellular processes in both roots and shoots, enabling plants to adapt efficiently to biotic and abiotic stimuli.

Our findings reveal that KIN10 overexpression results in constitutively active stress responses, particularly in shoots. Such transcriptomic landscape is often linked with stunted growth and increased resistance to pathogen attack, as an outcome of growth-defence trade-off. Contrary, *kin10* displays impaired root growth. The responses of both lines are reflected in substantial transcriptional changes. *kin10* roots, display pathways altered known to be crucial for signal perception, cell wall modulation, intracellular trafficking, and kinase activity, whereas KIN10-OE shoots show constitutively activated responses towards biotic stimuli.

Plant growth and development rely on the precise coordination of environmental perception and internal signalling networks. Among the most critical environmental cues are biotic interactions, which require spatially and temporally regulated responses to ensure both growth and survival. Roots, in direct contact with the soil environment, are specialized for sensing and responding to microbial signals. Root hairs, as extensions of the root epidermis, play a vital role in nutrient uptake and interactions with both symbiotic and pathogenic microbes. KIN10 integrates energetic and environmental cues to modulate developmental and defence-related pathways. While KIN10’s role in general stress responses has been well documented, its organ-specific functions, particularly in roots and shoots, remain incompletely understood. Overexpression of KIN10 leads to reduced shoot growth, which has been attributed to a shift toward constitutive stress signalling. Yet, how this overactivation modulates ROS dynamics and immune readiness in aerial tissues is less well explored. In summary, our findings establish SnRK1, through the function of its catalytic subunit KIN10, as a pivotal integrator of growth and defence programs in *Arabidopsis thaliana*, highlighting its crucial role in mediating the balance between resource allocation and environmental responsiveness. Building on this foundation, future research should prioritize the identification of SnRK1 phosphorylation targets, explore the tissue-specific regulation of its activity, and further dissect its role in shaping biotic interactions. Additionally, unravelling how SnRK1 interacts with other key signalling networks, such as hormonal and ROS pathways, will deepen our understanding of the intricate regulatory circuits governing plant adaptation.

## 4. Materials and methods

### 4.1. Plant material and cultivation

Experiments were performed on *Arabidopsis thaliana Col-0* and *Ler-0* plants. Transgenic lines of *A. thaliana kin10* (*Col-0* background) and KIN10-OE (*Ler-0* background) were previously described. Seeds were surface sterilized in a 30 % bleach solution (SAVO©, Unilever, Czech Repulic) with a droplet of Tween 20 for 5 min, rinsed 4 times with distilled water and then stored for stratification at 4 °C for 2 nights.

For the experiments with seedlings, plants were cultivated in transparent 24-well plates, in 400 μL of liquid Murashige-Skoog medium with vitamins (pH 5.7) 3-5 seedlings per well, for 10 days at short-day conditions (10 h light / 14 h dark). At 7^th^ day of cultivation 200 μL of fresh medium was added into each well.

For the experiments with mature leaves, seeds were sown in Jiffy 7 peat pellets and plants were cultivated at 22 °C, 70 % relative humidity, under a short-day photoperiod (10 h light/14 h dark), at 100-130 μE m^-2^ s^-1^. Plants were watered with distilled water and used for experiments when 4-to 5-weeks-old.

For the root experiments, seeds were placed on 9 cm diameter Petri plates filled with half-strength Murashige-Skoog medium supplemented with 0.8% agar, 20 seeds per plate. Petri plates were placed vertically in the D-ROOT system (8) to allow root elongation in the dark and cultivated for 7 days at 22°C, 16 h light/8 h dark regime.

### 4.2. Transcription networks establishment (RNAseq)

Total RNA was isolated from approximately 50 mg of root tissues using RNeasy Plant Mini kit (Qiagen) and treated with DNA-Free kit (Thermo Fischer Scientific). RNA purity, concentration and integrity were evaluated on 0.8% agarose gels (v/w) and by the RNA Nano 6000 Assay Kit using Bioanalyzer instrument (Agilent Technologies).

#### RNA-Seq Workflow

For RNA-seq analysis, approximately 5 μg of RNA were submitted for the service procedure provided by Eurofins Genomics. The analysis resulted in at least 10 million 150 bps read pairs. Rough reads were quality-filtered using Rcorrector and Trim Galore scripts (9).Transcript abundances (transcripts per million – TPM) were determined using Salmon (10) with parameters --posBias, --seqBias, -- gcBias, --numBootstraps 30. Index was built from TAIR10 *Arabidopsis thaliana* cds dataset. Visualization, quality control of data analysis and determination of differentially expressed genes were determined using sleuth (version 0.29.0) package in R (11). Transcripts with q-value <= 0.05 and log2 fold change >=1 (upregulated) or <= -1 (down-regulated) were considered to be significantly differentially expressed.

Functional Insights: Enrichment analysis of DEGs highlighted key biological processes through Gene Ontology (GO) terms.

### 4.3. Root length and root hairs formation in response to bacterial flagellin and living bacteria

To assess the root growth inhibition in response to bacterial elicitor flagellin, seedlings cultivated vertically on agarized medium (4 days post germination) were transferred to plates containing 100 nM flg22 and cultivated for another 3 days before scanning (Epson Perfection V700 Photo, Suwa, Japan, at 600 dpi resolution).

For the quantification of the lateral roots, plates were observed under a stereo microscope (SteREO Discovery V8, Carl Zeiss GmbH, Jena, Germany) equipped with an AxioCam HRc camera. Images were analyzed with FiJi software. For the measurement of root hair length and density, seedlings were photographed under an ApoTome Zeiss microscope with a 5x objective at bright field settings. Images were imported into FiJi software (12) and root hair length was measured manually using a segmented line tool.

Root hair formation in response to *Pseudomonas syringae* pv. *tomato* DC3000 (*Pst*)(7).

### 4.4. ROS formation in leaves

Rapid ROS formation was assessed by luminol-peroxidase assay. Discs 4 mm diameter were placed in white 96-well plate 150 μL dH2O and incubated overnight at RT to overcome mechanical stress. Water was replaced by 200 μL of assay solution containing 50 mM Tris-HCl, pH 8.5, buffer solution containing 17 μg.mL-1 luminol (Sigma-Aldrich), 10 μg.mL-1 horseradish peroxidase and AGP (0.1 mg/mL, 0.2 mg/mL, 0.5 mg/mL) or 100 nM flg22. Relative luminescence was measured during a 1 h period at 2 min intervals. Every well was measured separately and data from 12 independent measurements were used in one set of experiments.

### 4.5. Resistance to bacterial infection

Bacteria (wild-type *Pst* DC3000 and effector-deficient mutated strain PtoΔvrPtoΔAvrPtoB*-*) were cultivated overnight on plates containing LB medium (tryptone 10 g/L, NaCl 10 g/L, yeast extract 5 g/L, pH=7.0) supplemented with 1.4% agar and 50 mg/L rifampicin. Four-week-old plants (three leaves at the similar developmental stage, middle age leaves: 8th-9th-10th leaves) were syringe-infiltrated with the suspension of *Pst* (OD_600_=0.0005 in 10 mM MgCl_2_). At 2 days post inoculation, 3 discs (6 mm in diameter) were sampled from inoculated leaves from each plant, pooled (one plant as one sample) and homogenized in 1 mL of 10 mM MgCl_2_, in a 2 mL Eppendorf tube, with 1 g of 1.3 mm silica beads using a FastPrep-24 instrument (MP Biomedicals, USA). The resulting homogenate was subjected to serial 10× dilutions and pipetted onto LB plates. Colonies were counted after 1–2 days of incubation at 26 °C. Six individual plants were used per treatment, bacterial load being expressed as log10(CFU*mm^-2^)(13).

## References

1. Van Leene J, Eeckhout D, Gadeyne A, Matthijs C, Han C, De Winne N, et al. Mapping of the plant SnRK1 kinase signalling network reveals a key regulatory role for the class II T6P synthase-like proteins. Nat Plants [Internet]. 2022;8(11):1245–61. Available from: 10.1038/s41477-022-01269-w

2. Retzer K, Weckwerth W. The TOR–Auxin Connection Upstream of Root Hair Growth. Plants [Internet]. 2021;10(1). Available from: https://www.mdpi.com/2223-7747/10/1/150

3. Nukarinen E, Nägele T, Pedrotti L, Wurzinger B, Mair A, Landgraf R, et al. Quantitative phosphoproteomics reveals the role of the AMPK plant ortholog SnRK1 as a metabolic master regulator under energy deprivation. Sci Rep [Internet]. 2016;6(1):31697. Available from: 10.1038/srep31697

4. Margalha L, Confraria A, Baena-González E. SnRK1 and TOR: modulating growth–defense trade-offs in plant stress responses. J Exp Bot [Internet]. 2019 Apr 15;70(8):2261–74. Available from: 10.1093/jxb/erz066

5. Nukarinen E, Nägele T, Pedrotti L, Wurzinger B, Mair A, Landgraf R, et al. Quantitative phosphoproteomics reveals the role of the AMPK plant ortholog SnRK1 as a metabolic master regulator under energy deprivation. Sci Rep [Internet]. 2016;6(1):31697. Available from: 10.1038/srep31697

6. Baena-González E, Rolland F, Thevelein JM, Sheen J. A central integrator of transcription networks in plant stress and energy signalling. Nature [Internet]. 2007 Aug;448(7156):938—942. Available from: 10.1038/nature06069

7. Pečenková T, Janda M, Ortmannová J, Hajná V, Stehlíková Z, Žárský V. Early Arabidopsis root hair growth stimulation by pathogenic strains of Pseudomonas syringae. Ann Bot [Internet]. 2017 Sep 1;120(3):437–46. Available from: 10.1093/aob/mcx073

8. Silva-Navas J, Moreno-Risueno MA, Manzano C, Pallero-Baena M, Navarro-Neila S, Téllez-Robledo B, et al. D-Root: a system for cultivating plants with the roots in darkness or under different light conditions. The Plant Journal [Internet]. 2015 Oct 1;84(1):244–55. Available from: 10.1111/tpj.12998

9. Song L, Florea L. Rcorrector: efficient and accurate error correction for Illumina RNA-seq reads. Gigascience [Internet]. 2015 Dec 1;4(1):s13742-015-0089-y. Available from: 10.1186/s13742-015-0089-y

10. Patro R, Duggal G, Love MI, Irizarry RA, Kingsford C. Salmon provides fast and bias-aware quantification of transcript expression. Nat Methods [Internet]. 2017;14(4):417–9. Available from: 10.1038/nmeth.4197

11. Pimentel H, Bray NL, Puente S, Melsted P, Pachter L. Differential analysis of RNA-seq incorporating quantification uncertainty. Nat Methods [Internet]. 2017;14(7):687–90. Available from: 10.1038/nmeth.4324

12. Schindelin J, Arganda-Carreras I, Frise E, Kaynig V, Longair M, Pietzsch T, et al. Fiji: an open-source platform for biological-image analysis. Nat Methods [Internet]. 2012;9(7):676–82. Available from: 10.1038/nmeth.2019

13. Kalachova T, Janda M, Šašek V, Ortmannová J, Nováková P, Dobrev IP, et al. Identification of salicylic acid-independent responses in an Arabidopsis phosphatidylinositol 4-kinase beta double mutant. Ann Bot [Internet]. 2020 Apr 25;125(5):775–84. Available from: 10.1093/aob/mcz112

